# GASOTRANSMITTER MODULATION OF HYPOGLOSSAL MOTONEURON ACTIVITY

**DOI:** 10.1101/2022.03.23.485481

**Authors:** Brigitte Browe, Ying-Jie Peng, Jayasri Nanduri, Nanduri R. Prabhakar, Alfredo J. Garcia

## Abstract

Obstructive sleep apnea (OSA) is characterized by sporadic collapse of the upper airway leading to periodic disruptions in breathing. Upper airway patency governed by genioglossal nerve activity originates from the hypoglossal motor nucleus. Mice with targeted deletion of the gene *Hmox2,* encoding the carbon monoxide (CO) producing enzyme, heme oxygenase-2 (HO-2), exhibit severe OSA, yet the contribution of central HO-2 dysregulation to the phenomenon is unknown. Using the rhythmic brainstem slice preparation, which contains the preBötzinger complex (preBötC) and the hypoglossal nucleus, we tested the hypothesis that central HO-2 dysregulation weakens hypoglossal motoneuron output. Disrupting HO-2 activity increased transmission failure as determined by the intermittent inability of the preBötC rhythm to trigger output from the hypoglossal nucleus. Failed transmission was associated with a reduced input-output relationship between the preBötC and the motor nucleus. These network phenomena were related to smaller inspiratory drive currents and reduced intrinsic excitability among hypoglossal neurons. In addition to HO-2, hypoglossal neurons also expressed the CO-regulated H_2_S producing enzyme cystathionine □-lyase (CSE). H_2_S abundance was higher in hypoglossal neurons of HO-2 null mice than wild-type controls. Disrupting CSE function normalized transmission in HO-2 null mice and an H_2_S donor mimicked the effects of HO-2 dysregulation. These findings demonstrate a hitherto uncharacterized modulation of hypoglossal activity through the interaction of HO-2 and CSE-derived H_2_S, and supports the perspective that centrally derived HO-2 activity plays an important role regulating upper airway control.

## Introduction

Obstructive sleep apnea (OSA) is a prevalent breathing disorder affecting nearly a billion people throughout the world^1, 2^. It is characterized by the periodic cessation of breathing during sleep due to sporadic upper airway collapse. When left untreated, OSA predisposes the individual to a variety of diseases including hypertension^3, 4^, diabetes^5, 6^, and cognitive decline^7, 8^. Multiple factors contribute to the genesis of OSA including compromised pharyngeal anatomy ^9, 10^, inadequate upper airway muscle function^11–13^, low arousal threshold^14^, and a hypersensitive chemoreflex (i.e., high loop gain ^15^).

Peng et al. recently reported that mice with deletion of the *Hmox 2* gene, which encodes the enzyme heme oxygenase 2 (HO-2), exhibit a high incidence of OSA^16^. OSA in HO-2 null mice was attributed in part to the increased loop gain arising from the heightened carotid body chemo reflex^16–19^. While HO-2 produces several bioactive products^20^, the enhanced carotid body chemo reflex and subsequent OSA phenotype is attributed to the loss of HO-2 dependent CO production^16^. However, we hypothesize that additional factors, such as changed activities in the central nervous system, may also contribute to the OSA phenotype observed in HO-2 null mice.

Sporadic loss of neuromuscular control over upper airway muscles is a key contributor to producing obstructive apneas^21, 22^. Predisposition to intermittent reductions in airway patency can originate from changes in CNS activity. Disrupting rhythmic excitation of the hypoglossal nucleus that drives genioglossal nerve activity increases the likelihood for the tongue to occlude the upper airway during inspiration. Such disruptions may involve changing the state-dependent balance between excitation and inhibition received by respiratory hypoglossal motor neurons^23^ and/or the direct modulation of intrinsic excitability of hypoglossal neurons^24–26^. It, however, is unknown whether impaired HO-2 signaling in the hypoglossal nucleus influences synaptic and/or intrinsic neuronal properties to alter output from the motor nucleus that may ultimately contribute to upper airway obstruction.

We tested this possibility using a combination of electrophysiological, genetic, and pharmacological approaches in the rhythmic medullary brainstem slice preparation. We found that dysregulated HO-2 activity in the hypoglossal nucleus acts through CSE-dependent H_2_S signaling to reduce motor neuron excitability. This in turn, diminishes the input-output relationship between the preBötC and hypoglossal nucleus, and increases the likelihood of transmission failure between the premotor rhythm and motor nucleus output. These observations indicate that hypoglossal HO-2 / CO and CSE / H_2_S activities interact as important modulators of hypoglossal output that potentially contributes to changed upper airway tone when dysregulated.

## Methods

### Study Approval

In accordance with National Institutes of Health guidelines, all animal protocols were performed with the approval of the Institute of Animal Care and Use Committee at The University of Chicago (ACUP 72486, ACUP 71811).

### Experimental Animals

Experiments were performed using neonatal (postnatal day 6 to postnatal day 12) wildtype mice (C57/BL6; Charles River), HO-2 null mice (from S. H. Snyder; The Johns Hopkins University), and HO-2:CSE double-null mice. HO-2:CSE double-null mice were created by crossing HO-2 null females with CSE null males (from R. Wang, Department of Biology, Laurentian University, Sudbury, ON, Canada). Tissues from both sexes were used. No sex-based differences were observed; therefore, all sexes were pooled for analysis. All litters were housed with their dam in ALAAC-approved facilities on a 12 hour / 12-hour light-dark cycle.

### Pharmacological Agents

Heme oxygenase activity was blocked using bath application of Chromium (III) Mesoporphyrin IX chloride (ChrMP459, 20μM; Frontiers Sciences, Newark DE). CORM-3 (20μM; Sigma-Aldrich St. Louis MO), a CO-donor, was bath applied following ChrMP459 application. NaHS (10μM to 100 μM; Sigma-Aldrich), a H_2_S donor, was bath applied. In all patch clamp experiments, fast synaptic glycinergic and GABAergic inhibition was blocked by bath application of strychnine (1μM; Sigma-Aldrich), and picrotoxin (50μM; Sigma-Aldrich), respectively. Inhibition of CSE production was accomplished by *in vivo* L-propargylglycine (L-PAG, 30mg/kg (Sigma-Aldrich) administered (*i.p.* injection) 2.5 to 3 hrs prior to preparation of the rhythmic brainstem slice preparation. Inhibition of potassium channels SK_Ca_ and ATP-sensitive potassium channel (K_ATP_) was via bath application of APAMIN (200 μM; Sigma-Aldrich) and Tolbutamide (100 μM; Sigma-Aldrich) respectively.

### Measurement of H_2_S Production

Coronal brainstem sections (300μm thick) were cut with a cryostat at –20°C. The hypoglossal nucleus and control brainstem regions were excised with a chilled micro-punch needle. Hypoglossal tissue from a single brainstem was not sufficient for effectively measuring H_2_S levels; therefore, we pooled micro punched tissue from two mice for each sample where H_2_S levels measured. H_2_S levels were determined as described previously^27^. Briefly, cell homogenates were prepared in 100□mM potassium phosphate buffer (pH 7.4). The enzyme reaction was carried out in sealed tubes. The assay mixture in a total volume of 500μL contained (in final concentration): 100□mM potassium phosphate buffer (pH 7.4), 800μM□L-cysteine, 80μM pyridoxal 5′-phosphate with or without L-PAG (20µM) and cell homogenate (20μg of protein), was incubated at 37°C for 1□hr. At the end of the reaction, alkaline zinc acetate (1% mass / volume; 250μL) and trichloroacetic acid (10% vol/vol) were sequentially added to trap H_2_S and stop the reaction, respectively. The zinc sulfide formed was reacted with acidic N,N-dimethyl-p-phenylenediamine sulfate (20μM) and ferric chloride (30μM) and the absorbance was measured at 670□nm using Shimadzu UV-VIS Spectrophotometer. L-PAG inhibitable H_2_S concentration was calculated from a standard curve and values are expressed as nanomoles of H_2_S formed per hour per mg of protein.

### Immunohistochemistry

Anaesthetized mice (urethane, 1.2g•kg^−1^ *i.p.*) were perfused transcardially with heparinized phosphate-buffered saline (PBS) for 20 min followed by 4% paraformaldehyde in PBS. Brainstems were harvested, post-fixed in 4% paraformaldehyde overnight, and cryoprotected in 30% sucrose/PBS at 4°C. Frozen tissues were serially sectioned at a thickness of 20μm (coronal section) and stored at –80°C. Sections were treated with 20% normal goat serum, 0.1% bovine serum albumin and 0.1% Triton X-100 in PBS for 30 min and incubated with primary antibodies against choline acetyltransferase (ChAT, 1:100; Millipore; #AB144P), HO-2 (1:200, Novus Biologicals; # NBP1-52849) and CSE (1:250; gift from Dr. Schenider) followed by Texas Red-conjugated goat anti-mouse IgG or FITC-conjugated goat anti-rabbit IgG (1:250; Molecular Probes). After rinsing with PBS, sections were mounted in Vecta shield containing DAPI (Vector Labs) and analyzed using a fluorescent microscope (Eclipse E600; Nikon).

### Brainstem Slice for Electrophysiology

The isolated rhythmic brainstem slice was prepared as previously described ^28^. Briefly, animals were euthanized by decapitation. Brainstems were rapidly dissected, isolated, and placed into ice cold artificial cerebral spinal fluid (aCSF) (composition in mM: 118 NaCl, 25 NaHCO_3_, 1 NaH_2_PO_4_, 1 MgCl_2_, 3 KCl, 30 Glucose, 1.5 CaCl_2_, pH=7.4) equilibrated with 95% O_2_, 5% CO_2_. The isolated brainstem was glued to an agar block (dorsal face to agar) with the rostral face up and submerged in aCSF equilibrated with carbogen. Serial cuts were made through the brainstem until the appearance of anatomical landmarks such as the narrowing of the fourth ventricle and the hypoglossal axons. The preBötC and XIIn was retained in a single transverse brainstem slice (thickness: 560 ± 40μm). The slice was transferred into the recording chamber (∼6mL volume) where it was continuously superfused with recirculating aCSF (flow rate: 12 -15mL/min). Prior to recording, extracellular KCl was raised to 8mM and the spontaneous rhythm was allowed to stabilize prior to the start of every experiment.

### Electrophysiology

Extracellular population activity was recorded with glass suction pipettes filled with aCSF. Pipettes were positioned over the ventral respiratory column containing the preBötC and over the medial dorsal column containing the hypoglossal nucleus. In some experiments, a third pipette was positioned between the preBötC and hypoglossal nucleus just lateral to the axon tract to record transmission through the premotor field containing intermediate premotor inspiratory neurons^29, 30^. Extracellular population activity was recorded with glass suction pipettes filled with aCSF^31^. The recorded signal was sampled at 5kHz, amplified 10,000X, with a lowpass filter of 10 kHz using an A-M instruments (A-M Systems, Sequim, WA, USA) extracellular amplifier. The signal was then rectified and integrated using Clampfit electronic filter. Recordings were stored on a computer for *posthoc* analysis.

All intracellular recordings were made using the Multiclamp 700B (Molecular Devices: https://www.moleculardevices.com/systems/conventional-patch-clamp/multiclamp-700b-microelectrode-amplifier). Acquisition and post hoc analyses were performed using the Axon pCLAMP10 software suite (Molecular Devices: www.moleculardevices.com/system/axon-conventional-patch-clamp/pclamp-11-software-suite).

Whole cell patch clamp recordings of hypoglossal motor neurons were obtained using the blind-patch technique with a sample frequency of 40 kHz. Recordings were made with unpolished patch electrodes, pulled from borosilicated glass pipettes with a capillary filament^31^. The electrodes had a resistance of 3.5-8 MΩ when filled with the whole cell patch clamp pipette solution containing (in mM): 140 K-gluc acid, 1 CaCl_2_, 10 EGTA, 2 MgCl_2_, 4 Na_2_-ATP, 10 HEPES. Patch clamp experiments were performed with a patch clamp amplifier (Multiclamp 700B, Molecular Devices, Sunnyvale, CA, USA), and the software program pCLAMP 10.0 (Molecular Devices). Neurons located at least two to three cell layers (about 75-250μm) rostral from the caudal surface of the slice were recorded. The liquid junction potential was calculated to be -12mV and was subtracted from the membrane potential. The series resistance was compensated and corrected throughout each experiment. In voltage clamp experiments, membrane potential was held at -60mV. Current clamp experiments used a holding potential between 0 and -100pA to establish the baseline resting membrane potential between -55 and - 70mV. In some cases, we determined rheobases using a ramp protocol in our current clamp recordings. This ramp protocol consisted of a hyperpolarizing step (-100pA) succeeded by the injection of a ramping depolarizing current (122pA/sec; peak current 600pA).

### Statistical Analyses

Unless otherwise explicitly stated elsewhere, each n value represents an individual animal that served as a biological replicate for a given measurement. Transmission was expressed as a percentage of the hypoglossal network bursts corresponding to the total network bursts from either the preBötC or the premotor field. Burst were considered corresponding if initial start time of bursts were within 500-750ms of each other (corresponding time was maximized until only one hypoglossal burst per preBötC was detected). Mean I/O and transmission values for each slice were calculated using a 120 s window. This window was taken at the end of each baseline or pharmacological agent phase (each phase duration = 600 sec). The input-output (I/O) ratio for each inspiratory event (defined by network a burst in preBötC) was calculated as previously described in ^31^. I/O ratios values for preBötC bursts without a corresponding XIIn burst were designated as 0. To illustrate the cycle-to-cycle input-output relationships between networks, heat maps of I/O ratio values were plotted for each slice included in the experiment. Each row represents sequential cycles from a single slice experiment. As the rhythmic frequency across preparations varied, the number of events (i.e., cycle number) in the 120 s analysis window also varied; therefore, either the total number of cycles or 25 consecutive cycles from in a given analysis window were plotted.

Statistics were performed using Origin 8 Pro (OriginLab, RRID:SCR_014212) or Prism 6 (GraphPad Software; RRID:SCR_015807). In cases where the distribution of data appeared normal, comparisons between two groups were conducted using either paired or unpaired two-tailed t-tests as appropriate. In cases, where the distribution of individual data points did not appear normal, the Wilcoxon match-paired signed rank test was performed. A one-way ANOVA was performed followed by *posthoc* Dunnett’s test comparing experimental groups to control when a comparison of three or more groups. In plots where the mean ± S.E.M. are presented, the mean and S.E.M. are overlaid on the individual results from the corresponding dataset. Differences were defined when the P-value was less than 0.05.

## Results

### Hypoglossal neurons express hemeoxygenase-2 (HO-2)

We first assessed whether hypoglossal neurons express HO-2. Hypoglossal neurons showed positive immunohistochemical expression for HO-2 as indicated by co-localization of HO-2 with Choline acetyl transferase (ChAT), an established marker of these neurons ^11^ (**Fig.1A**, n=3).

**Figure 1.**
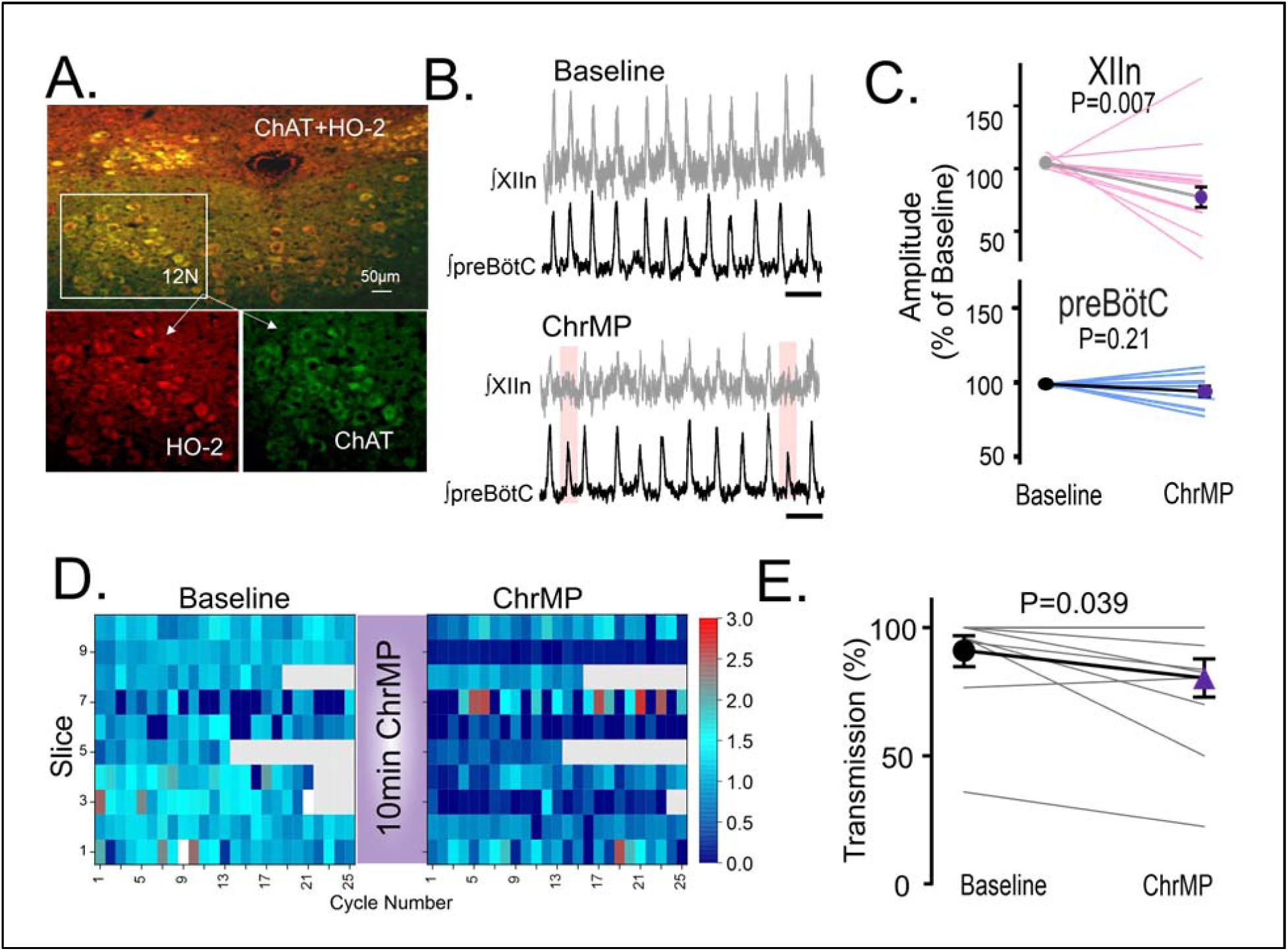
Disruption of hemeoxygenase-2 (HO-2) in hypoglossal motor neurons impairs inspiratory activity from the hypoglossal nucleus but not in the preBötC. **A.** HO-2 (red bottom left) is expressed in ChAT^+^ cells (green bottom right) of the hypoglossal nucleus (XIIn, combined top). **B-E**. Population recordings of rhythmic brain slices were recorded from ipsilateral preBotC and XIIn simultaneously, analyses were performed during baseline and during bath application of 20µM ChrMP459 (ChrMP). **B.** Integrated traces of network activity in spontaneously rhythmic brainstem slices (n=11) recorded from XIIn (grey) and preBötC (black) before (top) and after (bottom) bath application, failed transmission events are highlighted (pink boxes); scale bar: 5 sec. **C.** Percentage change of integrated burst amplitude from Baseline to ChrMP in XIIn (top) and the preBötC (bottom). Thin pink and blue lines illustrate integrated burst amplitude from individual slices. Symbols connected by a thick line illustrate mean integrated burst amplitude. **D.** Heat maps of consecutive cycle-to-cycle I/O ratios in Baseline and in ChrMP. Each row represents a single slice experiment during baseline and in ChrMP. Grey boxes indicate non-events in recordings from slower rhythms where less than 25 events (i.e., 25 cycles) occurred during the analysis window. **E.** Comparison of inspiratory drive transmission in XIIn between Baseline and ChrMP. Thin grey lines illustrate transmission from individual slices. Symbols connected by a thick line illustrate mean transmission.

### Disrupting HO-2 function impairs hypoglossal inspiratory activity

Extracellular field recordings in the rhythmic brainstem slice were simultaneously recorded from the preBötC and the corresponding motor output from the hypoglossal nucleus. Two approaches were employed to assess the role of HO-2: (1) using Cr(III) Mesoporphyrin IX chloride (ChrMP459, 20µM), an inhibitor of HO^32^; and (2) using brain slices from HO-2 null mice.

Representative extracellular field recordings from the preBötC and hypoglossal nucleus prior and during ChrMP459 exposure were shown in **Fig 1B** (n=11). While ChrMP459 suppressed hypoglossal burst amplitude (**Fig 1C**, *left*; n=11; Baseline: 99.57 ± 0.60%, ChrM459: 81.80 ± 10.80%, P=0.007), the HO inhibitor had no effect on preBötC burst amplitude (**Fig 1C**, *right*; n=11; Baseline: 99.93 ± 0.34%, ChrMP459: 93.23 ± 4.80%, P=0.21). ChrMP459 consistently reduced the cycle-to-cycle input-output relationship between preBötC and the motor nucleus as revealed by examining the cycle-to-cycle I/O across preparations (**Fig 1D**, Baseline: 1.00 ± 0.07 vs. ChrMP459: 0.55 ± 0.11; P=.006). In the extreme, altered input-output relationships between preBötC and the hypoglossal nucleus may increase the propensity for transmission failures as determined by the inability of preBötC activity to produce hypoglossal output at the network level^31^. Indeed, the reduced I/O ratio was associated with an increase in failed transmissions of the preBötC activity to output from the hypoglossal nucleus (**Fig 1E**, Baseline: 90.85 ± 5.80% vs. ChrMP459: 80.21 ± 7.40%, P=0.039). Together these findings suggest that HO inhibition causes a generalized weakening in the activity relationship between the rhythm generating network and hypoglossal motoneurons.

ChrMP459 is a pan HO inhibitor; however, it cannot distinguish activities between heme oxygenase isoforms. To assess the specific contribution of HO-2, we compared rhythmic activities in brain slices from wild type (n=9) and HO-2 null (n=7) mice (**Fig 2A**). Larger cycle-to-cycle I/O ratios were observed in wild type slices as compared to HO-2 null slices (**Fig 2B**: wild type: 0.99 ± 0.04 vs. HO-2 null: 0.76 ± 0.11, P=0.045). Similarly, transmission of preBötC activity to the hypoglossal nucleus was greater in wild type than in HO-2 null slices (**Fig 2** **C,** wild type: 96.53 ± 1.76% vs. HO-2 null: 62.55 ± 7.93%, P=0.0006). These findings established that genetic elimination of HO-2 produces a similar phenomenon to that of pan HO inhibition indicating that loss of HO-2 activity alone is sufficient for impairing transmission from the preBötC to the hypoglossal nucleus. Given these similarities and the limited availability of HO-2 null mice, several of the following studies were performed using the ChrMP459 in rhythmic wild type brainstem slices.

**Figure 2.**
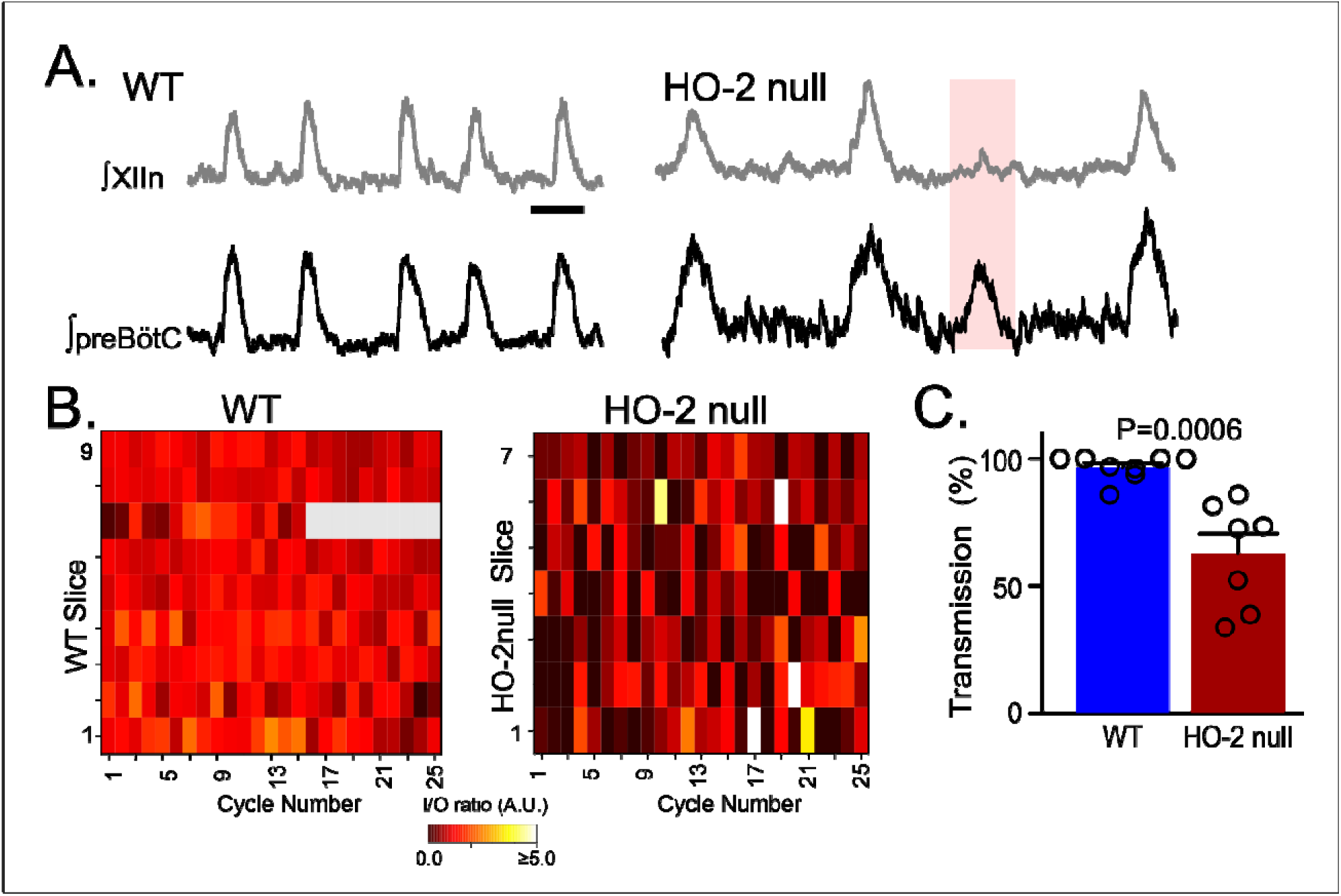
Genetic deletion of HO-2 reduces the I/O relationship between preBötC and the hypoglossal nucleus and uncouples of motor output from inspiratory rhythmogenesis. **A.** Representative integrated traces of network burst activity in the preBötC and XIIn in rhythmic slices from wildtype (WT) (left, n=9) and HO-2 null (HO-2^-/-^; right, n=7) mice; scale bar: 2 sec. **B.** Heat maps of cycle-to-cycle I/O ratios in WT (left) and HO-2^-/-^ (right) slices. Grey boxes indicate non-events in recordings from slower rhythms where less than 25 events occurred during the analysis window. **C.** Transmission of inspiratory activity from preBötC to XIIn in slices from WT vs HO-2^-/-^. Symbols illustrate transmission from individual slices.

### HO inhibitor does not affect premotor neuron activity

Intermediary premotor neurons relay drive from the preBötC to the hypoglossal nucleus^29, 30^. Therefore, it was possible that HO inhibition impaired transmission of drive from the preBötC by perturbing activity from intermediary premotor neurons. To address this possibility, triple extracellular recordings (n=5) were made from the preBötC, the field of the ipsilateral premotor neurons, and the hypoglossal nucleus. Baseline transmission from the preBötC to the premotor field and to the hypoglossal nucleus was reliable and consistent (**Fig. 3A**, *middle panel*). However, ChrMP459 disrupted activity in the hypoglossal nucleus despite unaltered activities in either the preBötC or the intermediate premotor field (**Fig 3A**, *right panel*). Indeed, while neither transmission failures nor the cycle-to-cycle I/O ratio from the preBötC to the premotor field was affected by ChrMP459 (**Fig 3B**: *left;* I/O: Baseline: 1.16 ± 0.09 vs ChrMP 1.16 ± 0.14, P=0.31; *right;* Transmission: Baseline: 100.0 ± 0.0% vs ChrMP 86.35 ± 11.81%, P=0.312), the HO inhibitor reduced the transmission of activity and the cycle-to-cycle I/O ratio between the premotor field and the hypoglossal nucleus (**Fig 3C**: *left,* I/O: Baseline: 1.12±.17 vs ChrMP 0.40±0.09, P=0.04; *right,* Transmission: Baseline 89.57±1.3% vs ChrMP 57.62±13.69%, P=0.04). These results suggested that dysregulated HO-2 modulates is a postsynaptic phenomenon in the hypoglossal nucleus.

**Figure 3.**
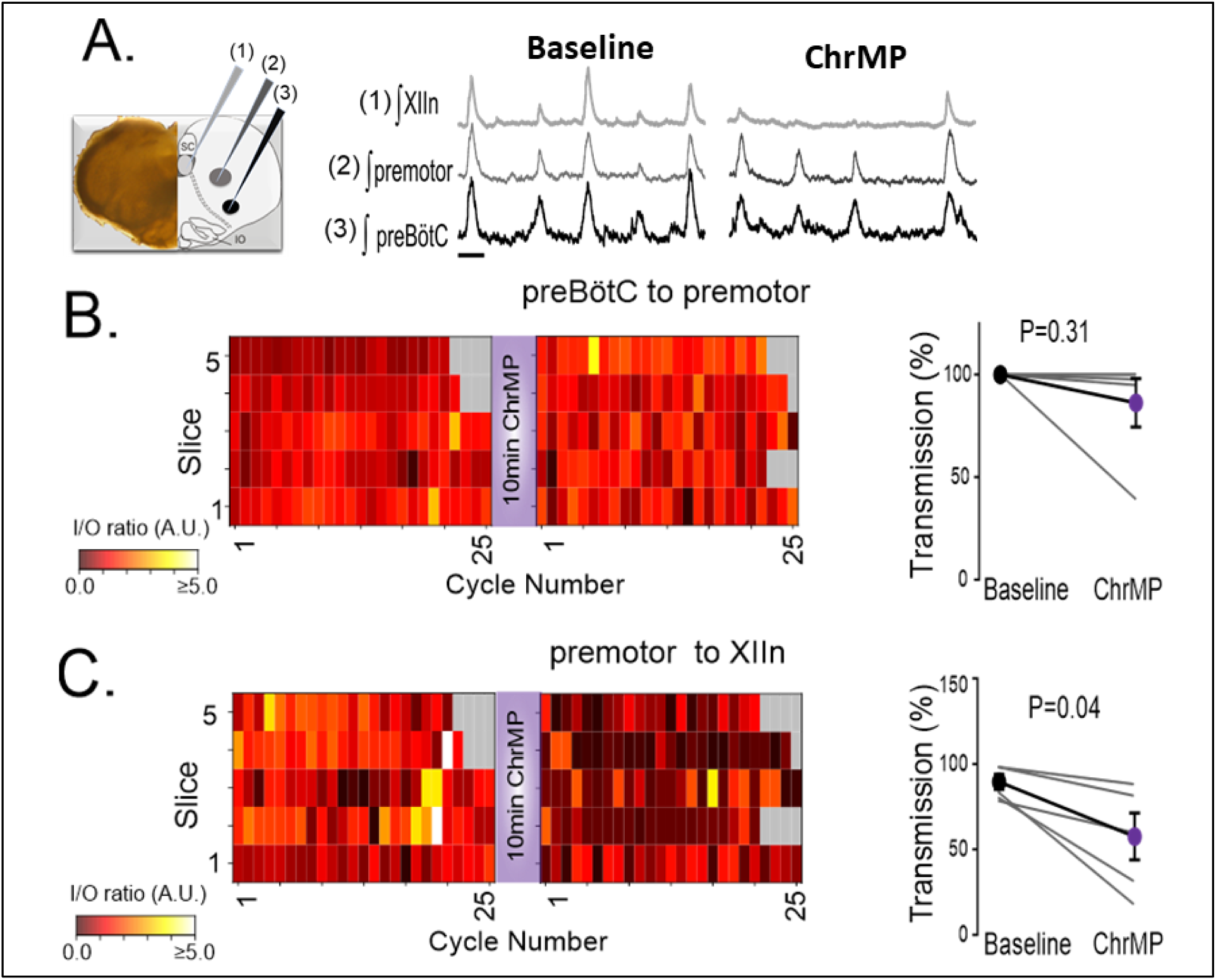
While ChrMP459 does not change transmission from the preBötC to the premotor area, ChrMP459 increases transmission failure from the premotor area to the hypoglossal nucleus. **A.** Diagram of medullary brain slice illustrating relative electrode placement for simultaneous triple extracellular recordings (n=5) from the XIIn (light grey, 1), premotor field (dark grey, 2), and preBötC (black, 3). Corresponding representative traces of integrated network activity in Baseline (left) and in 20μM ChrMP (right); scale bar: 1 sec. **B.** Heat maps of the cycle to cycle I/O ratio (left) and transmission (right) between preBötC and the premotor field. **C.** Heat maps of the cycle to cycle I/O ratio (left) and transmission (right) between the premotor field and XIIn.

### HO inhibition suppresses inspiratory drive currents and reduces excitability in hypoglossal neurons

To examine the effect of HO inhibition on postsynaptic activity of the hypoglossal neurons, we performed patch clamp recordings from a total of 27 wild type hypoglossal neurons exposed to ChrMP459. These hypoglossal neurons were disinhibited from fast inhibition (50μM picrotoxin and 1μM strychnine) allowing us to focus on inspiratory-related fast glutamatergic drive. 19 of the 27 hypoglossal neurons received excitatory synaptic drive in-phase with the preBötC (i.e., inspiratory hypoglossal neurons). Peak inspiratory drive currents were reduced in ChrMP459 (**Fig 4A**, n=19, Baseline: -142.9 ± 22.82 pA vs. ChrMP459: -95.31 ± 21.79 pA, P=0.004). Reduced drive coincided with hypoglossal neurons generating fewer action potentials per preBötC burst in ChrMP459 (**Fig 4B**, n=17, Baseline: 14.68 ±2.23 action potentials per burst vs. ChrMP459: 6.79± 1.54 action potentials per burst, P<0.0001). Furthermore, as determined by the injection of a depolarizing ramp current into hypoglossal neurons, the HO inhibitor increased rheobase among inspiratory hypoglossal neurons (**Fig 4C**, n=19, Baseline: 167.5 ± 35.83 pA vs. ChrMP459: 338.0 ± 82.50 pA; P=0.008) yet decreased rheobase in non-inspiratory hypoglossal neurons (i.e., neurons not receiving drive during preBötC activity; **Supplemental Fig 1**, n=8, Baseline: 280.5±56.43 pA vs 228.2±47.96pA, P=0.017).

**Figure 4.**
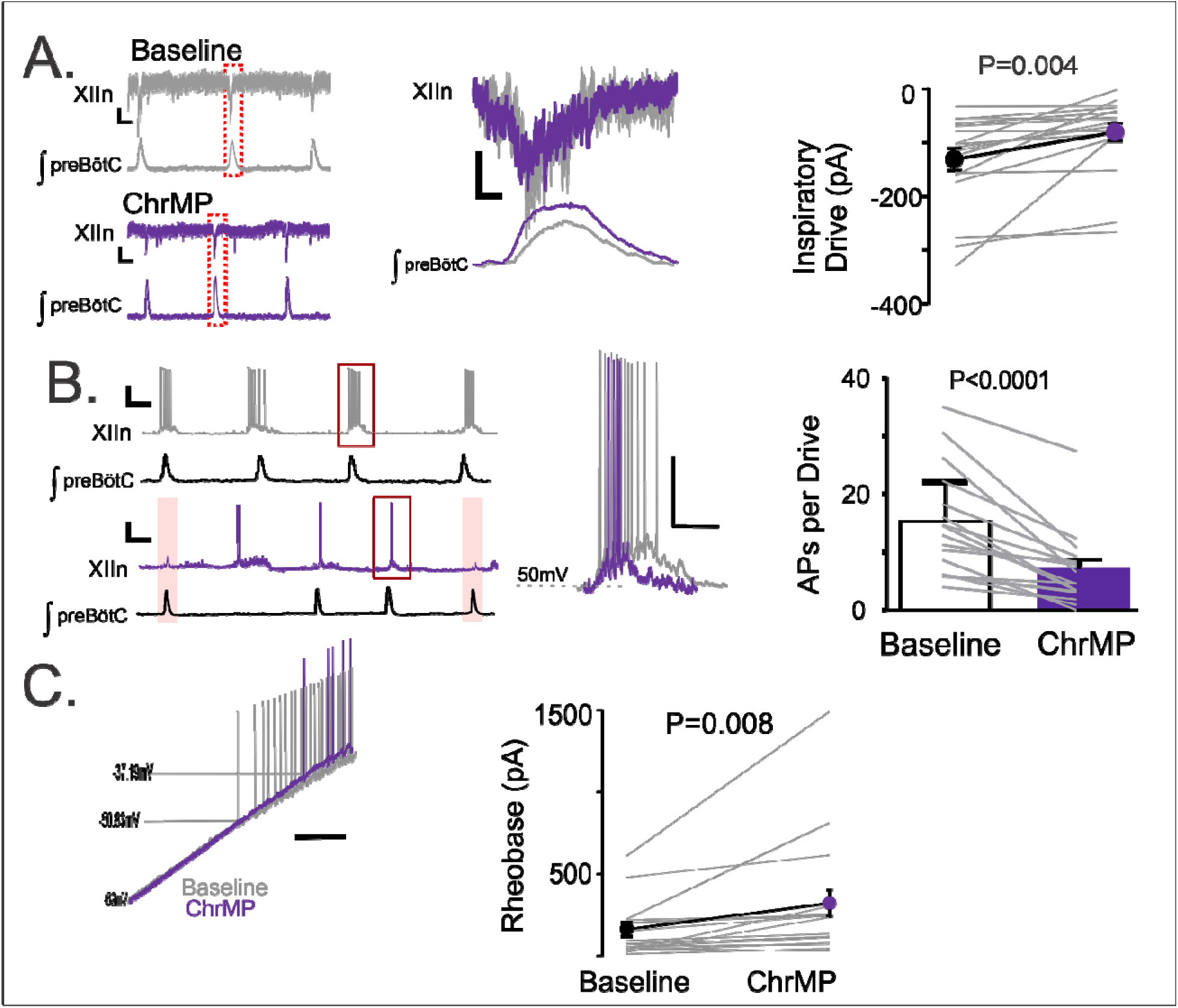
Heme oxygenase inhibition reduces inspiratory drive currents in hypoglossal neurons. Whole cell patch clamp recordings were made from hypoglossal neurons in rhythmic brain slices while simultaneously recording ipsilateral preBötC activity in Baseline and in ChrMP. Neurons were disinhibited from fast synaptic inhibition using 50μM PTX and 1μM Strychnine. **A.** (left) Representative voltage clamp recordings from a XIIn neuron (V_holding_= -60mV) aligned with corresponding integrated network activity from preBötC before (Baseline, top, grey) and after 20 μM ChrMP (bottom, purple). Scale bar: 1 sec x 10pA. (middle) Magnification of highlighted (red dotted box) drive currents from Baseline (grey) and ChrMP (purple). Scale bar: 1000 msec x 10pA. (right) Comparison of XIIn inspiratory drive current magnitude in Baseline and ChrMP (n=19). Thin grey lines illustrate individual neuron response. Symbols connected by a thick line illustrate mean drive current. **B.** (left) Representative current clamp recordings from a XIIn neuron spontaneously active with the preBötC network rhythm in Baseline (top, grey) and 20 μM ChrMP (bottom, purple); skipped transmission of action potentials in ChrMP are highlighted (pink). Scale bar 2sec x 20mV. (middle) Magnification of highlighted neuronal activity (red box in left). Scale bars: 100msec x 25mV. (right) Number of action potentials generated per inspiratory burst in Baseline and in ChrMP (n=17). Thin grey lines illustrate individual neuron response. **C.** (left) Representative trace of current clamp recording in response to ramp current injection during Baseline (grey trace) and in ChrMP (purple trace); scale bar: 500 msec. (right) Comparison of rheobase in inspiratory XIIn neurons during Baseline and in ChrMP (n=19). Thin grey lines illustrate individual neuron response.

### Elevated levels of H_2_S are observed in the hypoglossal nucleus of HO-2 null mice

We next sought to determine the mechanism(s) by which inhibition of HO-2 affect hypoglossal neuron activity. Earlier studies^33, 34^ have reported that HO-2 is a negative regulator of CSE-dependent H2S production. To test this possibility, we first examined whether the hypoglossal neurons express CSE. Brainstem sections from the wild type hypoglossal nucleus revealed CSE hypoglossal tissue punches from wild type and HO-2 null mice. Relative to the wild type hypoglossal nucleus, H_2_S is expressed in ChAT-positive hypoglossal neurons (**Fig 5A**, n=3). We then determined H_2_S abundance in hypoglossal tissue punches from wild type and HO-2 null mice. Relative to the wild type hypoglossal nucleus (**Fig 5B** *blue*; n=6, 60.58 ± 6.37 nmol • mg^-1^ • h^-1^), H_2_S abundance was greater in the hypoglossal nucleus of HO-2 null mice (**Fig 5B** *red*; n=6, 144.12 ± 8.29 nmol • mg^-1^ • h^-1^), but not different from the inferior olive brainstem region of HO-2 null mice (**Fig 5B** *grey*; n=4, 56.10 ± 2.88 nmol • mg^-1^ • h^-1^). These findings suggest that the hypoglossal nucleus expresses CSE and HO-2 negatively regulates H_2_S production in the hypoglossal nucleus.

**Figure 5.**
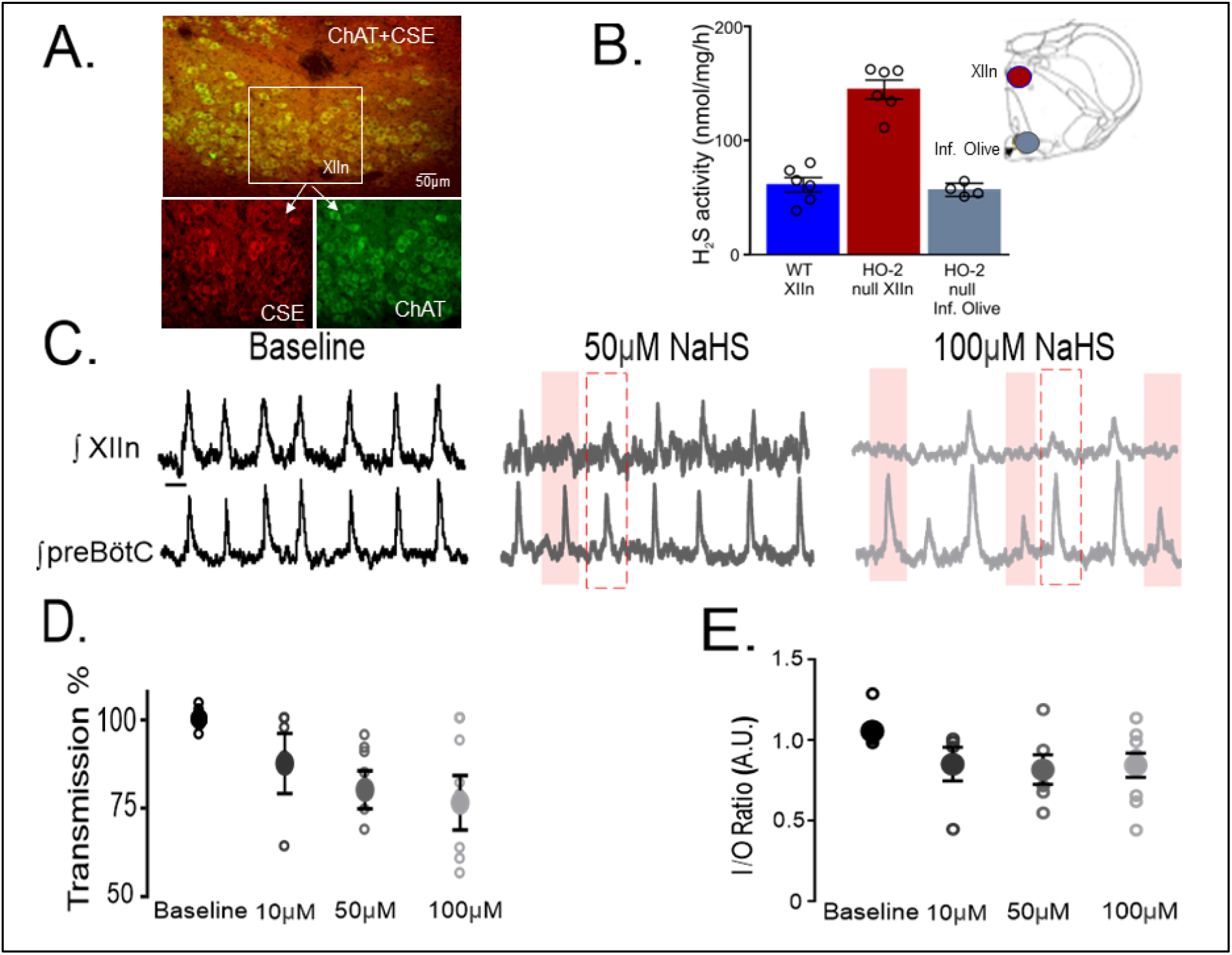
CSE-dependent H_2_S is produced in the hypoglossal nucleus and exogenous NaHS uncouples hypoglossal nucleus activity from the preBötC. A. **A.** CSE (red, bottom left) is expressed in ChAT^+^ neurons (green, bottom right) in the XIIn (overlay, top). **B.** CSE-dependent H_2_S generation in homogenates from WT and HO-2^-/-^. Homogenates were prepared from tissue punches from the XIIn (red area in slice diagram) and inferior olive (grey area in slice diagram) at bregma between -7.20mm and -7.90mm. (WT: XIIn n=6; HO-2^-/-^: XIIn n=6, inferior olive n=4). Note: Each n value reported in B represents a sample consisting of the anatomical region pooled from two animals. **C.** Integrated traces from XIIn (top) and preBötC (bottom) during baseline (black), and in response to the H_2_S donor, NaHS, at 50 μM (dark grey) and 100 μM (light grey). After NaHS application, XIIn but not preBötC burst amplitude were diminished (red dashed box) and in some cases, preBötC drive failed to produce activity in th XIIn (pink boxes). **D.** Comparison of transmission from preBötC to XIIn after NaHS application at 10μM, 50μM and 100μM. **E.** I/O ratios for each NaHS concentration. (Baseline: n=9; 10μM n=5; 50μM n=6; 100μM n=9).

### H_2_S mediates impaired transmission of inspiratory drive caused by disrupted HO-2 function

If the impaired transmission of inspiratory drive to the hypoglossal nucleus by disrupted HO-2 function involves CSE-derived H_2_S then: 1) a H_2_S donor should mimic the effects of disrupted HO-2 activity; 2) CO administration should improve the input-output relationship in respiratory slices from HO-2 null mice and ChrM459 application; and 3) CSE blockade should restore the transmission from the preBötC to the hypoglossal nucleus. The following experiments tested these possibilities.

Wild type brainstem slices exhibited a nearly 1:1 ratio of transmission of inspiratory activity from preBötC to hypoglossal neurons (**Fig 5C**, *left*). Application of NaHS, a H_2_S donor reduced transmission from preBötC to hypoglossal (**Fig. 5C**, *middle, right*) in a dose-dependent manner (**Fig. 5D**; 0μM NaHS: n=9, 100 ± 0.73%; 10μM NaHS: n=5, 90.14 ± 6.08%; 50μM NaHS: n=6, 84.0 ± 3.91%; 100μM NaHS: n=9, 81.2 ± 5.83%), which coincided with a reduction in I/O ratio by NaHS (**Fig. 5E**; 0 μM NaHS: 1.055 ± 0.028; 10μM NaHS: 0.85 ± 0.09; 50μM NaHS: 0.82 ±0.08; 100μM NaHS: 0.84 ± 0.07). These findings demonstrated increased H_2_S abundance reduces hypoglossal neuronal activity consistent with findings using either ChrM459 or HO-2 null mice. Thus, the stability of inspiratory transmission from the preBötC to hypoglossal nucleus appears to be negatively affected either by increasing H_2_S abundance or by disrupting HO-2 function.

CO produced by HO-2 is known to inhibit CSE-dependent H_2_S production by HO-2 ^33, 34^. Therefore, we sought to assess how CORM-3 (20μM), a pharmacological CO donor, impacted activity in ChrMP459-treated wild type rhythmic slices (n= 3, **Fig 6A**) and rhythmic slices from HO-2 null mice (n=4). Dysregulated HO-2 activity, caused by either pharmacological (ChrMP459) or genetic (HO-2 null mice) manipulation, is improved by CORM-3 as indicated by larger I/O ratios (**Fig 6B**, dysregulated HO-2: 0.71 ± 0.04 vs. CORM-3: 1.03 ± 0.08, P=0.009) and improved transmission (**Fig 6C**, dysregulated HO-2: 79.88 ± 4.69% vs. CORM-3: 93.39 ± 2.93%, P=0.036). As these findings suggested the absence of HO-2 dependent CO production is a key factor driving transmission failure in the rhythmic slice preparation, we next determined the involvement of CSE.

**Figure 6.**
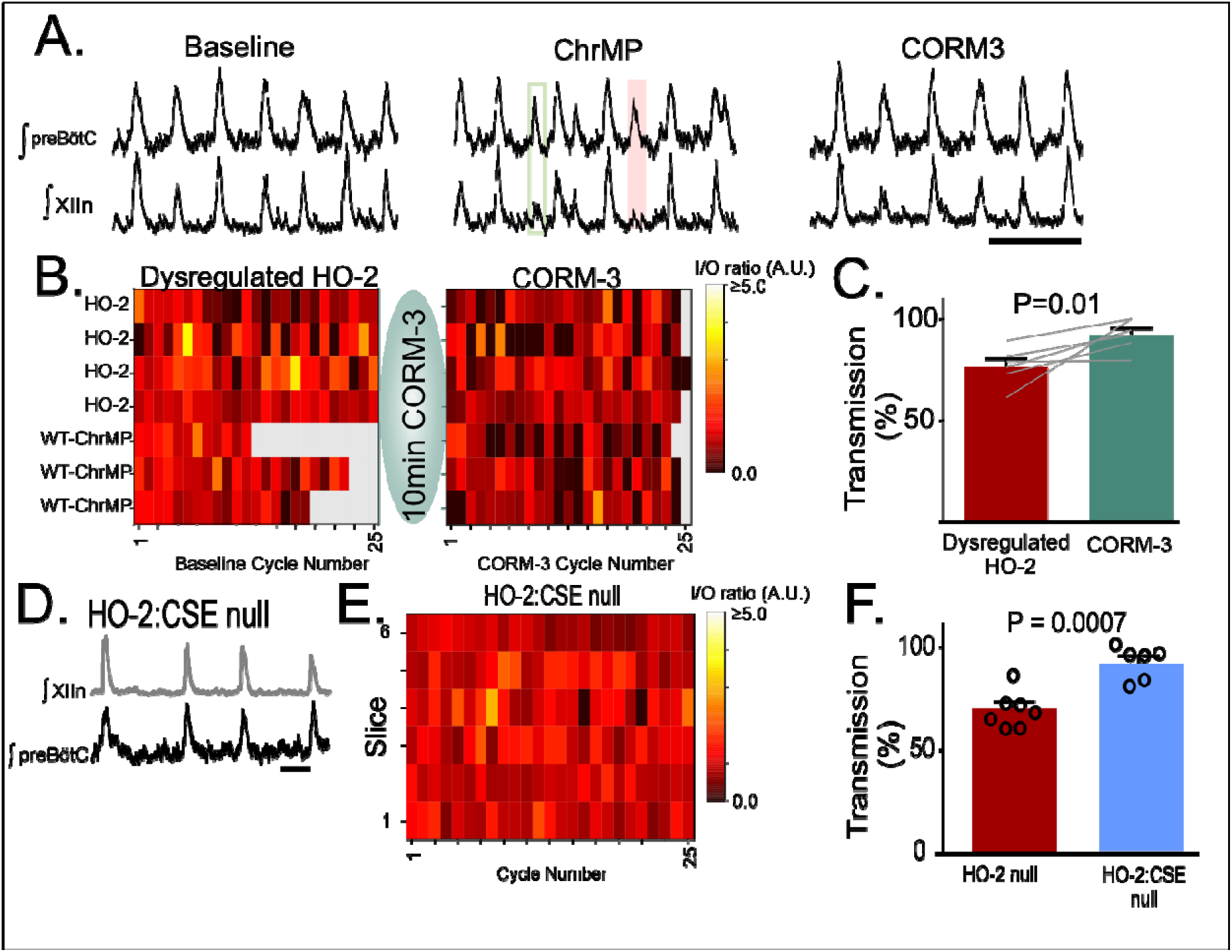
HO-dependent transmission failures can be recovered with CO-donor CORM-3 and are not present in HO-2:CSE null transmission. **A.** Representative extracellularrecording from preBotC and XIIn in WT slices during Baseline (left), in ChrMP alone (middle), and in ChrMP + 20uM CORM-3 (CORM3, right). **B.** Heat maps of cycle-to-cycle I/O ratios during dysregulated HO-2 (n=7: n=4 HO-2 null and n=3 WT-ChrMP) before and after CORM-3 application. **C.** Transmission from preBötC to XIIn from dysregulated HO-2 slices before and after bath application of CORM-3. **D.** Representative extracellular recordings from preBötC and XIIn in slices from HO-2:CSE null (HO-2:CSE^-/-^); scale bar 2 sec. **E.** Heat map of cycle-to-cycle I/O ratio from preBötC to XIIn in HO-2:CSE^-/-^. The I/O ratio from HO-2:CSE^-/-^ is greater than I/O ratios from HO-2^-/-^ (P=0.003). **F.** Comparison of transmission from preBötC to XIIn in HO-2^-/-^ and HO-2:CSE^-/-^ (n=6). HO-2^-/-^ data used for comparisons in E and F were originally shown in Figure 2.

Inspiratory activity in the brainstem slice from HO-2:CSE double null mice appeared to be stable and consistent (**Fig 6D**). Quantification of simultaneous extracellular field recordings of preBötC activity and hypoglossal nucleus revealed a larger I/O ratio (**Fig 6E**, HO-2:CSE: 1.04 ± 0.02, n=6; P=0.003) and near absence of transmission failures (**Fig 6F**, HO-2:CSE: 91.6 ± 0.02%; P=0.0007) when compared to activity in HO-2 null slices.

Given the observations using HO-2:CSE null mice, we next sought to determine whether acute blockade of CSE could restore transmission relationships between the preBötC and the hypoglossal nucleus in the HO-2 null slice. *In vivo* L-PAG treatment improved transmission of preBötC activity to the hypoglossal nucleus in the rhythmic slice (**Fig. 7A**, n=6) as indicated by larger cycle-to-cycle I/O ratios (**Fig. 7B**, L-PAG = 1.01± 0.03, P=0.008) and greater transmission rates (**Fig. 7C**, L-PAG 96.31 ± 2.62%, P<0.0001) when compared to the respective metrics from untreated HO-2 null mice. Intermittent transmission failure was also evident in patch clamp recordings from untreated HO-2 null hypoglossal neurons (**Fig 7D**, *left* shaded cycles) but not in HO-2 null hypoglossal neurons treated with L-PAG (**Fig. 7D**, *right*). These reduced transmission events correlated with smaller individual inspiratory drive currents in HO-2 null hypoglossal neurons when compared to corresponding inspiratory drive currents from L-PAG treated HO-2 mice (**Fig. 7D-E**, HO-2: -36.71 ± 2.14 pA vs. L-PAG -194.3 ± 82.73 pA, P=0.0007). Together, these experiments implicated the involvement of CSE-dependent H2S signaling with the effects of disrupted HO-2 signaling in hypoglossal neurons.

**Figure 7.**
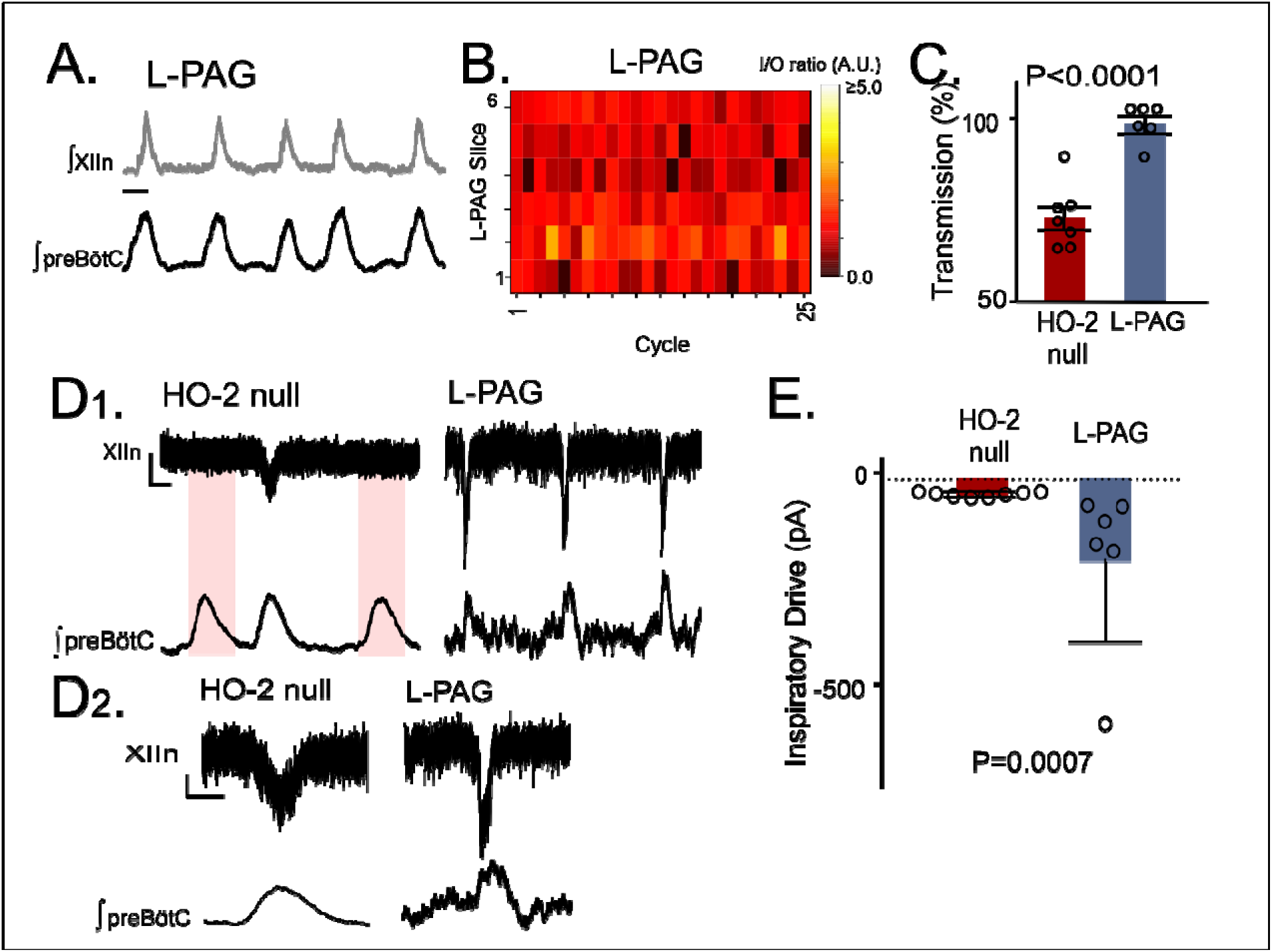
Transmission failures in HO-2 null mice are rescued with CSE inhibitor L-propargylglycine. **A.** Representative recordings of integrated activity of the preBötC (bottom) and XIIn (top) in slices from HO-2 null mice treated with L-PAG (30mg/kg; L-PAG, n=5). Scale bar 2 sec. **B.** Heat map of cycle-to-cycle I/O ratio in rhythmic slices from HO-2 null mice treated with L-PAG (n=6). **C.** Comparison of transmission between HO-2^-/-^ (replotted from Fig 2) and L-PAG slices (n=6). **D.** Representative voltage clamp recordings of inspiratory drive currents received by hypoglossal neurons from HO-2^-/-^ (n=7) and L-PAG (n=6). Neurons were disinhibited from fast synaptic inhibition using 50μM PTX and 1μM Strychnine. Scale bar 10 msec x 20pA. Skipped transmission between preBötC (bottom) the XIIn neuron (top) occurs in untreated HO-2^-/-^ (highlighted by pink boxes) but not in neurons from L-PAG. **D1.** Magnified representative drive potentials from HO-2^-/-^ and L-PAG (from highlighted by red-outlined boxes in **D**). scale bars: 100 msec x 10pA. **E.** Quantification of drive currents received by XIIn motoneurons from HO-2^-/-^ mice (n=8) produce smaller drive potentials when compared to L-PAG (n=6).

### Blockade of small conductance calcium-activated potassium channel (SK_Ca_) activity restores changes induced by HO-dysregulation in hypoglossal activity

As our experiments implicated the involvement of H_2_S signaling, we next sought to determine how H_2_S sensitive ion channels may contribute to impairing hypoglossal neuron activity caused by HO-dysregulation. H_2_S has been shown to enhance activity of several different potassium channels, including SK_Ca_ and ATP-sensitive potassium channel (K_ATP_) activities ^35^. As both SK_Ca_ and K_ATP_ are important in the regulation of hypoglossal neuron excitability ^36, 37^, we examined how blocking these channels affected hypoglossal activity during HO-dysregulation. At the network level, administration of the selective SK_Ca_ inhibitor, apamin (200μM), increased the excitability of the hypoglossal neurons treated with ChrMP459. This enhanced activity exceeded the original baseline activity (i.e., prior to ChrMP459 administration) causing ectopic bursting in the hypoglossal nucleus (**Supplemental Figure 2A**) making analysis of population transmission and I/O ratios unreliable. Therefore, we proceeded to resolve the effects of apamin on the influence of ChrMP459 at the level of individual hypoglossal inspiratory motor neurons. While in some hypoglossal neurons exposed to ChrMP459 apamin substantially increased drive currents (>100pA; **FIG 8A1**), in others, apamin modestly increased the drive current (<100pA; **FIG 8A2**). Despite this variability, apamin increased inspiratory drive currents received by ChrMP459 treated hypoglossal neurons (**FIG 8A3,** n= 6, ChrMP459: -85.77± 38.54 pA vs. apamin: -219.97± 97.76 pA, P=0.031). Apamin also enhanced the number of action potential generated per preBötC burst during ChrMP459 (**FIG 8B**, n=8, ChrMP-459: 12.57 ± 3.7 action potential per burst vs. apamin 26.05 ± 6.87 action potential per burst, P=0.016). This was consistent with the ability of apamin to reduce rheobase in ChrMP459 treated inspiratory hypoglossal neurons (**FIG 8B**; n=7, ChrMP459: 532.0 ± 186.5 pA vs. Apamin: 307.09 ± 80.9 pA, P=0.016).

**Figure 8.**
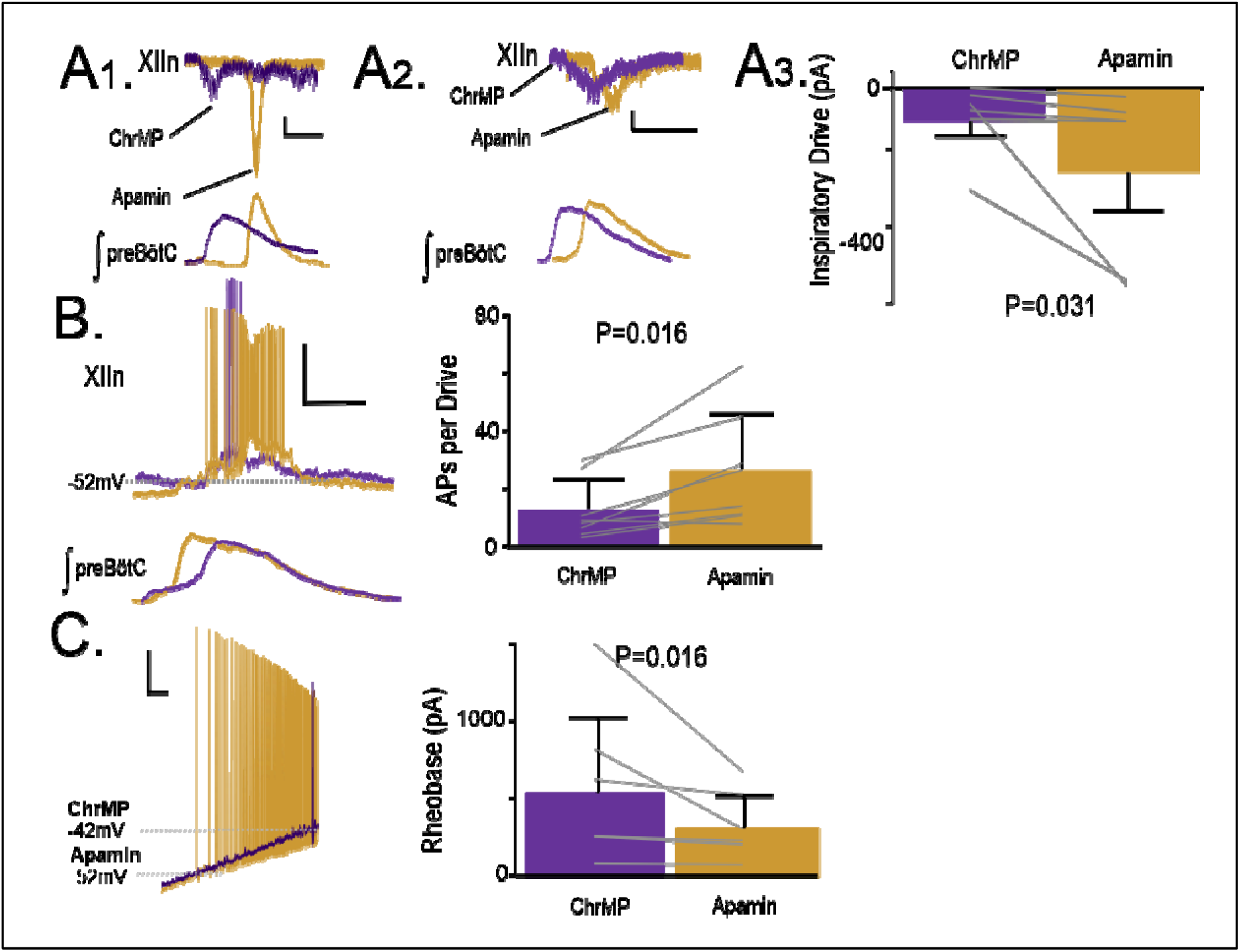
Apamin improves network input-output relationship between hypoglossal nucleus and reverses changes to intrinsic and synaptic excitability caused by HO dysregulation. **A1** Representative inspiratory drive current received by hypoglossal neurons in ChrMP (purple) and in Apamin (200 μM, gold) where apamin increased the drive current >100 pA. Scale bars: 1 s x 50 pA. **A2.** Representative inspiratory drive current hypoglossal neuron in ChrMP (purple) and in Apamin (gold) where apamin increased the drive current by less than <100 pA Scale bars: 1 s x 50 pA. **A3.** Comparison of inspiratory drive currents from hypoglossal neurons exposed to ChrMP (200 μM) then treated with Apamin (n=6). The effect of ChrMP on baseline disinhibited drive current for each of these neurons were reported in Fig 4A. **B**. (left) Representative trace of bursting of a hypoglossal neuron resulting from preBötC drive during ChrMP (purple) and in Apamin (gold) (n=8). Scale bars: 20 mV x 500 msec. (right) Comparison of action potentials generated per preBötC burst during ChrMP and Apamin (n=8). The effect of ChrMP on baseline action potential generated per preBötC burst for each of these neurons were reported in Fig 4B. **C.** (left) Representative traces of current clamp recordings in response to ramp current injection in ChrMP (purple) and in Apamin (gold). Scale bar: 500msec x 10mV). (right) Rheobase comparison from inspiratory XIIn neurons during ChrMP and Apamin (n=7). The effect of ChrMP on baseline rheobase for each of these neurons were reported in Fig 4C.

To determine how blockade of K_ATP_ impacted hypoglossal activity during ChrMP459, we used the K_ATP_ channel blocker, tolbutamide (100μM). Tolbutamide did not induce ectopic bursting in the hypoglossal nucleus during ChrMP459 (**Supplemental >Fig 2B**, n=5). Furthermore, tolbutamide (100μM), did not improve the rate of transmission of preBötC activity (**Supplemental Fig 2C**, ChrMP459: 69.24±6.0% tolbutamide: 71.23 ± 6.8%, P=0.76) but did increase the I/O ratio (**Supplemental Fig 2C** **right**, ChrMP459: 0.68 ± 0.05 tolbutamide: 0.87 ± 0.05, P=0.037). Tolbutamide neither enhanced inspiratory drive currents in ChrMP459 (**Supplemental Fig 2D**, n=4, ChrMP459: -70.51 ± 27.49 pA vs. tolbutamide: -83.06 ± 36.29 pA, P=0.37) nor increased the number of action potentials per preBötC burst in ChrMP459 (n=4, ChrMP459: 7.063±2.08 action potential per burst vs. tolbutamide: 9.37±3.18 action potential per burst, P=0.125). Moreover, tolbutamide did not affect rheobase of inspiratory hypoglossal neurons treated with ChrMP459 (**Supplemental Fig 2D**, n=4; ChrMP459: 221.01 ± 74.8 pA vs. tolbutamide: 180.4 ± 68.63 pA, P=0.218). Thus, these results suggested that apamin could enhance activity of hypoglossal neurons during HO-inhibition; whereas, the efficacy of tolbutamide to impact activity during HO-inhibition was limited.

## Discussion

Our study reveals a previously uncharacterized neuromodulatory interaction between HO-2 and CSE-derived H_2_S regulating activity from the hypoglossal nucleus. Our electrophysiological studies revealed that dysregulated HO-2 activity impacted intrinsic properties of hypoglossal neurons receiving drive from the preBötC. HO-2 dysregulation impaired transmission originating from the rhythm generating preBötC and the motor nucleus, as evidenced by the cycle-to-cycle reductions in the input-output relationship and the intermittent failures for preBötC activity to generate corresponding motor pool output, which mitigated blocking CSE activity and mimicked using a H_2_S donor. The increased propensity of transmission failure in response to dysregulated HO-2 activity and involvement of CSE / H_2_S are reminiscent of the OSA phenotypes observed in HO-2 null mice^16^. Thus, our study demonstrates the potential importance of centrally derived interactions between HO-2 and CSE activity in regulating motor neuron output important for maintaining upper airway patency.

Both a pharmacological inhibitor of HO and genetic elimination of Hmox-2 impaired transmission of neural drive from preBötC to the hypoglossal nucleus. Mice treated with intermittent hypoxia (IH) patterned after blood O_2_ profiles seen during OSA also show failed transmission related to a changed input-output relationship between preBötC and hypoglossal nucleus^31, 38^. While IH-induced phenomena are associated with alterations in preBötC activity, the HO inhibitor neither impacted inspiratory rhythmogenesis from the preBötC (**Fig. 1C**) nor perturbed transmission of neuronal drive to intermediate premotor neurons suggesting that changed interneuron neurophysiology was not a major contributing factor to the impaired transmission in the hypoglossal nucleus.

At the level of the hypoglossal nucleus, we documented that HO-2 is expressed among ChAT-positive cells of the hypoglossal nucleus. Although we did not resolve expression among hypoglossal motor neurons innervating upper airway muscles, such as genioglossal neurons, our electrophysiological studies demonstrated that ChrMP459 had divergent effects on non-inspiratory and inspiratory hypoglossal neurons. HO inhibition increased rheobase among non-inspiratory neurons; whereas, in inspiratory hypoglossal neurons, it decreased the magnitude of drive currents, increased rheobase, and diminished the number of action potentials generated during preBötC bursting. Thus, while our studies did not resolve how hypoglossal neurons projecting to different tongue muscles are impacted by ChrMP459, our findings indicate that HO-dysregulation can differentially impact hypoglossal neurons that receive drive from the preBötC.

The observed postsynaptic phenomena among hypoglossal motor neurons receiving inspiratory drive could contribute to the occurrence of obstructive apneas by increasing the probability for intermittent reductions in upper airway tone from inspiratory drive by hypoglossal motor neurons. Such loss of upper airway muscle tone can obstruct airflow even when inspiratory activity is successfully produced at in other inspiratory muscles, such as the diaphragm. Indeed, this phenotype has been documented in HO-2 null mice^16^. However, further resolution is needed to understand how dysregulation of HO-2 may impact hypoglossal neurons innervating different muscle groups and to determine extent of the contribution from central HO-2 dysregulation to loss of upper airway tone and airway collapse *in vivo*.

In HO-2 null mouse, the incidence of OSA is absent with co-inhibition of CSE^17^, which is consistent with reports that CO generated by HO-2 inhibits CSE-dependent H_2_S production^33, 34^. After documenting CSE expression in hypoglossal neurons and demonstrating an increased abundance of H_2_S in the hypoglossal nucleus of HO-2 null mice, we demonstrated that a CO donor improves transmission and the input-output relationship between the preBötC and the hypoglossal nucleus in HO-2 null slices. Furthermore, using a H_2_S donor also increased transmission failures and reduced the I/O ratio similar to perturbing HO-2 activity. Endogenous H_2_S activity could originate from other H_2_S producing enzymes, such as cystathionone β-synthase (CBS) that is expressed primarily in astrocytes throughout the CNS^39^. However, CBS inhibition appears to have limited impact on inspiratory activity from the hypoglossal nucleus^40^.

In contrast, our genetic and pharmacological manipulations to ablate/diminish CSE activity improved transmission and the input-output relationship between inspiratory drive and the hypoglossal nucleus in HO-2 null mice. Together these findings implicated the interaction between HO-2 / CO and CSE / H_2_S within the hypoglossal nucleus to regulate its output.

How might enhanced H_2_S signaling reduce drive currents, and excitability of hypoglossal neurons? While it is possible that H_2_S may impact presynaptic release of glutamate and/or postsynaptic receptor activity of hypoglossal neurons, the ChrMP459 mediated increase in rheobase among inspiratory hypoglossal neurons implicated the involvement of a non-synaptic conductance(s) downstream of H_2_S based signaling caused by perturbations in HO-2 activity. H_2_S can enhance both K_ATP_ and SK_Ca_ activities^35^. In the hypoglossal neurons, K_ATP_ is dynamically active causing periodic adjustment of neuronal excitability^36^ while SK_Ca_ also regulates excitability and firing properties of hypoglossal neurons^37^. In ChrMP459, tolbutamide had a limited effect normalizing transmission as it improved the I/O ratio between preBötC and the hypoglossal motor nucleus, but did not reduce transmission failure. At the neuronal level, tolbutamide did not increase drive currents nor did it enhance intrinsic excitability of hypoglossal neurons in ChrMP459. These results indicated that blockade of K_ATP_ during HO-2 dysregulation was limited in countering the effects on HO-2 dysregulation in the hypoglossal nucleus. On the other hand, apamin normalized drive currents and increased excitability of inspiratory hypoglossal neurons in ChrMP459 demonstrating that blockade of SK_Ca_ sufficiently mitigates many aspects of HO-2 dysregulation in hypoglossal neurons that receive drive from the preBötC.

In conclusion, our study provides proof-of-concept for the existence of a central mechanism by which loss of HO-2 leads to reduced upper airway tone by enhancing H_2_S activity in the hypoglossal nucleus. This mechanism appears to involve antagonistic interactions between HO-2 and CSE activities to regulate excitability of hypoglossal neurons and is localized in the neurons themselves. Although OSA in HO-2 null mice has attributed to increased “loop-gain” arising from the hypersensitive carotid body reflex^16^, our findings indicate the potential involvement of a disrupted interaction between HO-2 / CO and CSE / H_2_S in hypoglossal motor neurons that contribute could to the sporadic loss of upper airway tone observed in OSA.

## Acknowledgements

**Funding Sources:** This work was supported by NIH P01 HL144454 (NRP), NIH R01NS107421 (AJG), NIH R01HL163965 (AJG) and NIH R01DA057767 (AJG).

---

**Supplemental Figure 1:**
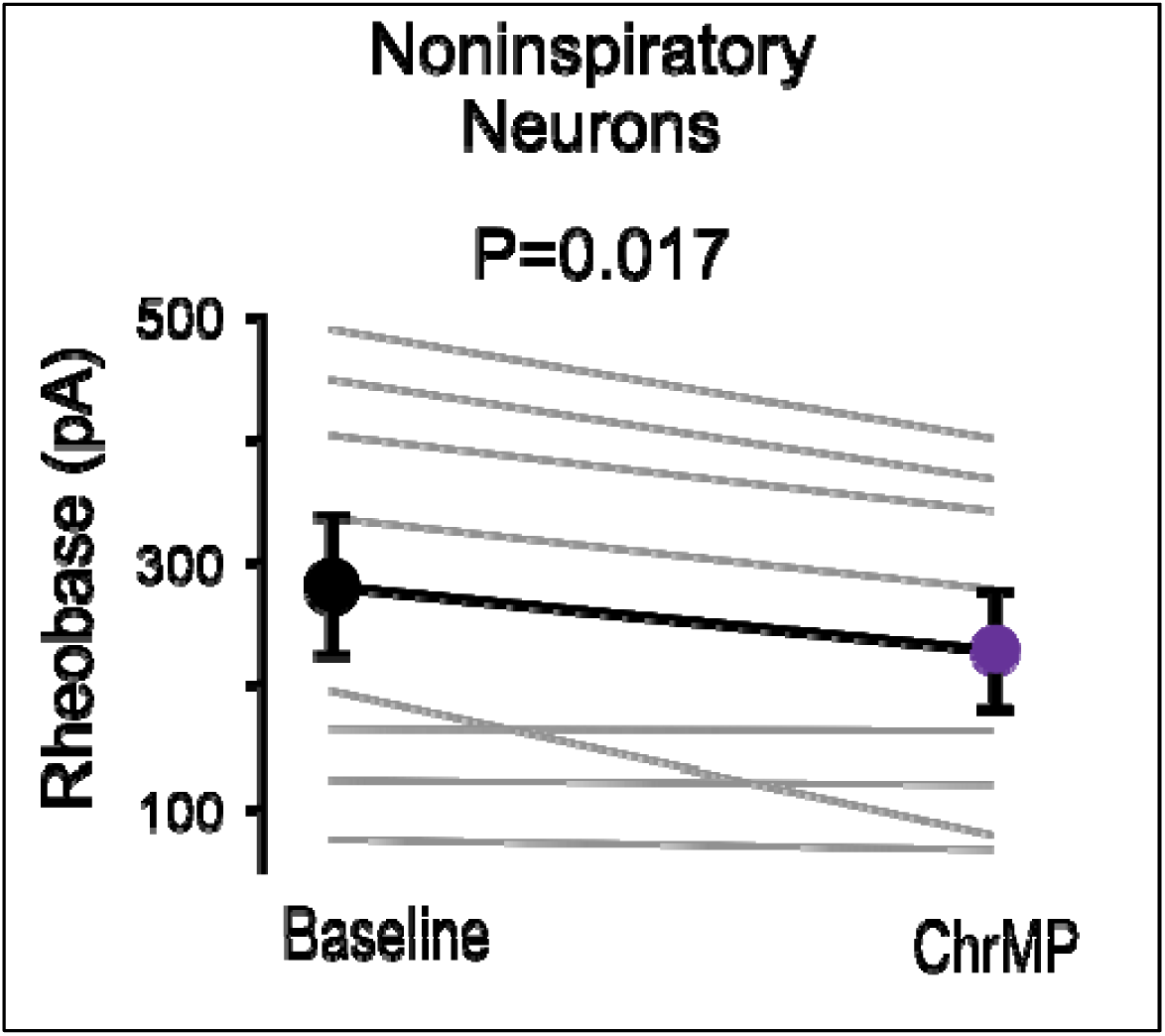
ChrMP459 decreases rheobase of non-inspiratory hypoglossal neurons. Comparison of rheobase from non-inspiratory hypoglossal neurons in baseline and ChrMP (n=8). Non-inspiratory hypoglossal neurons were defined as neurons that did not receive synaptic drive in-phase with a preBötC burst. Neurons were disinhibited from fast synaptic inhibition using 50μM PTX and 1μM Strychnine.

**Supplemental Figure 2:**
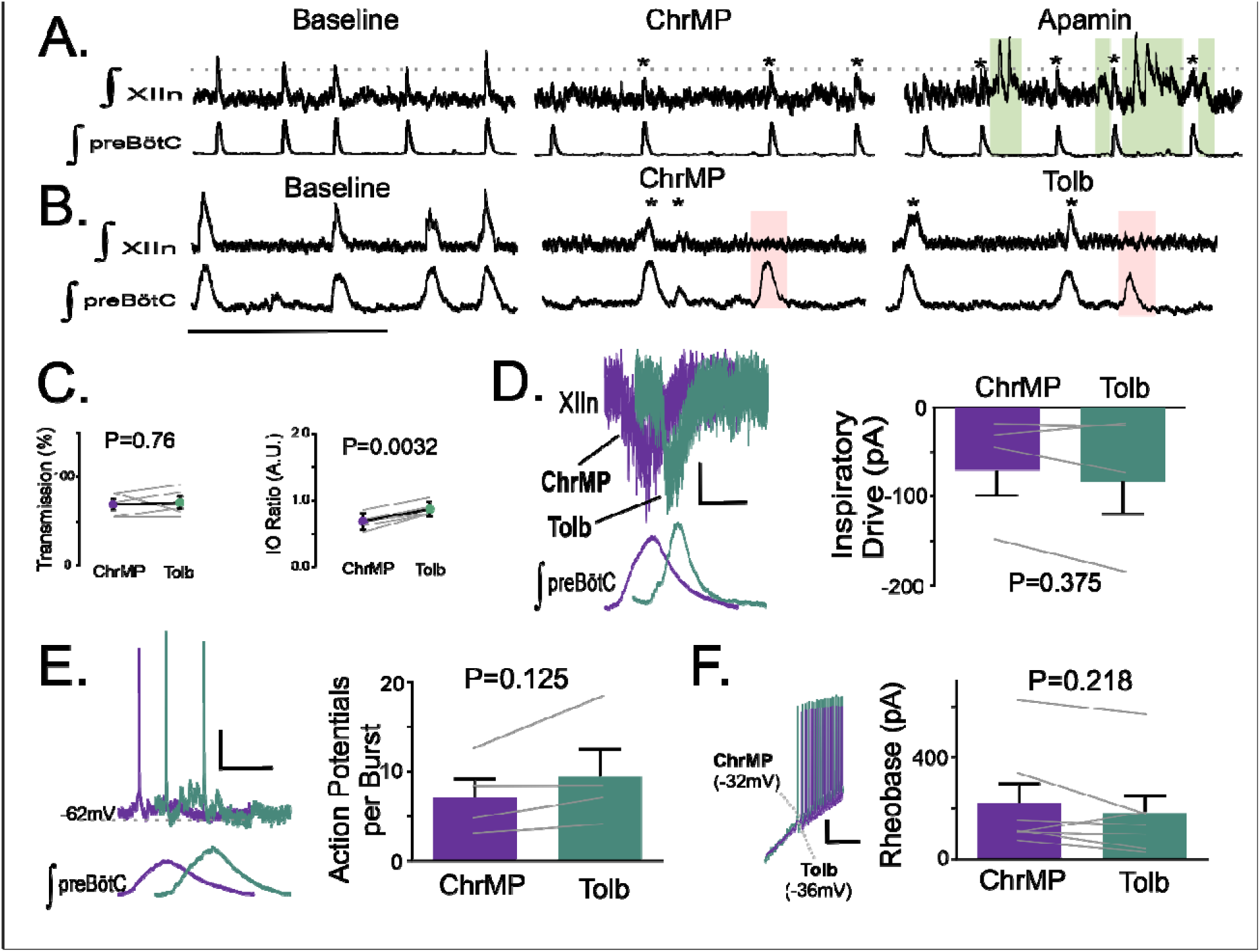
Apamin produce ectopic hypoglossal network activity during ChrMP459 and tolbutamide has limited effects on network and inspiratory hypoglossal neurons activity during ChrMP459. **A.** Representative traces of integrated network activity of the preBötC (top) and XIIn (bottom) in slices from WT mice Baseline, ChrMP, and in 200 μM Apamin. Asterisks (*) indicate detected hypoglossal output in phase with the preBötC. Green box indicates ectopic network bursting. Scale bar 10 sec. Due to ectopic bursting within and around rhythmic burst transmissions, analysis of apamin at the network level could not be accurately detected. **B.** Representative traces of integrated network activity of the preBötC (top) and XIIn (bottom) in slices from WT mice Baseline, ChrMP, and in tolbutamide (200 μM, Tolb). Asterisks (*) indicate detected hypoglossal output in phase with the preBötC. Pink box indicates cycles where preBötC drive failed to produce activity in the XIIn. **C.** Comparisons of Transmission (left) and I/O ratio (right) in ChrMP459 and in Tolbutamide (n=5). **D.** (top) Representative traces of inspiratory drive currents in ChrMP (purple) and in Tolb (green). Scale bar 1 sec x 50 pA. (bottom) Comparison disinhibited inspiratory drive currents in ChrMP and Tolb (n=4). The effect of ChrMP on baseline drive current for each of these neurons were reported in Fig 4B. **E**. (left) Representative trace of bursting of a hypoglossal neuron resulting from preBötC drive during ChrMP (purple) and in Tolb (green) (n=4). Scale bars: 10 mV x 500 msec. (right) Comparison of action potentials generated per preBötC burst during ChrMP and Tolb (n=4). The effect of ChrMP on baseline action potentials generated per preBötC burst for each of these neurons were reported in Fig 4C. **F.** (left) Representative traces of current clamp recordings in response to ramp current injection in ChrMP (purple) and in Tolb (green). Scale bar: 1 sec x 10mV). (right) Rheobase comparison from inspiratory hypoglossal neurons during ChrMP and Tolb (n=7). The effect of ChrMP on rheobase for each of these neurons were reported in Fig 4C.

